# A systematic review of climate-change driven range shifts in mosquito vectors

**DOI:** 10.1101/2025.03.25.645279

**Authors:** Kelsey Lyberger, Anna Rose Robinson, Lisa Couper, Isabel Delwel, Caroline Glidden, Crystal Qian, Aja Burslem, Faith Fernandez, Benjamen Gao, Gabriella Garcia, Julio Gomez, Caspar Griffin, Stephanie Jackson, Annalisa King, Olivia Manes, Andrew Song, Edward Tran, Erin A. Mordecai

## Abstract

As global temperatures rise, concerns about shifting mosquito ranges—and accompanying changes in the transmission of malaria, dengue, and other diseases—are mounting. However, systematic evidence for climate-driven changes in mosquito ranges remains limited. We conducted a systematic review of studies documenting expansions or contractions in medically important mosquito species. In total, 178 studies on six continents identified range expansions in 118 mosquito species. While over a third of these studies cited warming as a driver, fewer than 10% performed statistical tests of the role of climate. Instead, most expansions were linked to human-aided dispersal (e.g., trade, travel), land-use changes, and urbanization. Although several studies reported poleward or upward expansions consistent with climate warming, none demonstrated warm-edge contractions driven by rising temperatures, which are theoretically predicted in some settings. Rather than expanding into newly suitable areas, many expansions appear to be filling preexisting thermally suitable habitats. Our findings highlight the need for long-term mosquito monitoring, rigorous climate-attribution methods, and better documentation of confounding factors like land-use change and vector control efforts to disentangle climate-driven changes from other anthropogenic factors.

## Introduction

As anthropogenic carbon emissions continue to accelerate climate change, pervasive impacts are emerging on ecosystems, species distributions, and human health (IPCC AR6 WGII). In particular, climate change is causing escalating health threats due to exposure to climate extremes (e.g., heat waves and more intense storms), wildfires and their resulting smoke and toxic pollution, malnutrition and food insecurity, and climate-sensitive infectious diseases (Rocklöv et al. 2020, Rocklöv and Dubrow 2020, Watts et al. 2021, Childs et al. 2022, Burke et al. 2023, Franklinos et al. 2019). Although changes in vector-transmitted diseases were among some of the earliest hypothesized effects discussed in the literature on biological effects of climate change (Martens et al. 1997, Craig et al 1999, Rogers and Randolph 2006, Lafferty 2009, Parham and Michael 2010), evidence of climate-driven changes in vector-borne disease transmission remains limited and mostly consists of future projections (e.g., Ryan et al. 2015, Lippi et al. 2019, Ryan et al. 2021, Ryan et al. 2023, Semenza et al. 2023, but see Carlson et al. 2023a). Drawing on ecological theory and empirical work showing that organisms respond nonlinearly to temperature, with both lower and upper thermal limits for life history traits, Lafferty (2009) argued that climate change would not drive *expansions* of infectious diseases but range *shifts*, characterized by expansions at cool-edge limits and contractions at warm-edge limits. Yet, the extent to which these predicted range shifts have been occurring remains unclear as we do not yet have systematic scientific evidence on the degree to which observed expansions or contractions in vectors and infectious diseases are linked to climate change.

As ectotherms, the vectors of infectious diseases, including mosquitoes, ticks, flies, fleas, and other arthropods, are sensitive to temperature because it affects their vital rates, including development, survival, and reproduction (Angilletta 2009, Dell et al. 2011, Rogers and Randolph 2006). For mosquitoes, in particular, evidence of nonlinear effects of temperature on life history traits, with performance inhibited beyond both lower and upper thermal limits, is clear from both laboratory experiments and field data on disease transmission (Mordecai et al. 2019). Species distribution model studies, which aim to characterize the probability of species’ occurrence as a function of environmental variables using occurrence data from the field, consistently find temperature to be important in delimiting the species ranges of mosquitoes (e.g., Brady et al. 2014, Cunze et al. 2016, Messina et al. 2019, Lippi et al. 2023, Athni et al. 2024). Thus, theory predicts that, all else being equal, mosquito species should expand at their cooler range limits and contract at warmer range limits with climate change; however, other limiting factors such as habitat and host availability can also constrain range limits beyond physiological thermal limits (Lippi et al. 2023, Franklinos et al. 2019).

Although expansions and shifts in mosquito-borne diseases like malaria, dengue, West Nile, Zika, yellow fever, Japanese encephalitis, and others pose a major public health threat (Rocklöv and Dubrow 2020), the extent to which climate-driven mosquito range expansions and shifts are already occurring due to climate warming remains an important scientific evidence gap. One study of *Anopheles* malaria vectors in Africa found that species ranges have shifted an average of 4.7 km polewards and 6.5 m upwards in elevation per year over the last century, consistent with hypothesized effects of climate change (Carlson et al. 2023a). Likewise, mosquito range expansions into high elevation regions above 2000 m have been documented in Nepal for *Anopheles*, *Aedes*, and *Culex* vectors of malaria, dengue, and other arboviruses, respectively, where outbreaks of mosquito-borne diseases have also expanded dramatically (Dhimal et al. 2014a, Dhimal et al. 2014b, Dhimal et al. 2015, as summarized in IPCC AR6 WGII Ch 2). However, in many cases, range expansions of mosquitoes and mosquito-borne diseases have occurred in concert with land use change and socioeconomic factors that are concurrent with climate change, making it difficult to isolate the causal effects of climate change (Franklinos et al. 2019).

Beyond a few compelling case studies, major evidence gaps remain in whether mosquito range expansions are consistently occurring across species and geographic regions, whether they are attributable to climate change *per se*, and whether climate-driven range contractions are occurring at warm-edge limits. These knowledge gaps impede public health preparedness for emerging mosquito-borne disease outbreaks. In this systematic review we seek to address these gaps by: 1) synthesizing existing literature on mosquito range shifts, 2)determining the extent to which current literature attributes range shifts to climate change, 3) outlining alternative drivers of species range expansions, 4) synthesizing the speed of mosquito range expansions and its consequences for disease transmission, and 5) outlining the key open questions and future directions for understanding climate-driven vector range shifts. This review contributes to a more comprehensive understanding of current knowledge on the connections between climate, mosquito ecology, and public health, crucial for future prevention and intervention strategies and for climate policy.

## Methods

Following best practices for systematic reviews, we pre-registered our literature search methods on OSF: https://osf.io/w4gmu. We conducted a systematic review of studies related to changes in mosquito ranges. Searches were conducted in Pubmed, Scopus, Web of Science (Biosis, CABI, Zoological Record), Google Scholar, BioRxiv, and SciELO in April 2023 using the following search terms: ("range contraction*"[tw] OR "range expansion*"[tw] OR "range shift*"[tw] OR "northward expansion*"[tw] OR "elevation shift*"[tw] OR "elevational shift*"[tw] OR "altitude shift*"[tw] OR "altitudinal shift*"[tw] OR "first occurrence"[tw] OR "shifting distribution*"[tw] OR "distribution shift*"[tw] OR "expanding distribution*"[tw]) OR "distribution expan*"[tw] OR "contracting distribution*"[tw] OR "distribution contract*"[tw] OR "Animal Distribution"[Mesh]) AND ("Mosquito*"[tw] OR "Aedes"[tw] OR "Anopheles"[tw] OR "Culex"[tw] OR "Culicidae"[Mesh] OR "Mosquito Vectors"[Mesh]) AND ("English"[Language]). This search yielded 2356 studies after we removed duplicate records (see Fig. S1 for a flow chart of how studies were processed using the software platform Covidence in this systematic review).

Studies that were selected for inclusion contained primary evidence of range expansions or contractions for a medically relevant mosquito vector species (including but not limited to those in the genera *Aedes*, *Anopheles*, and *Culex*) and/or the pathogen(s) it transmits. Acceptable evidence of a range expansion was defined as new reports of a vector collected or recorded in the wild in a location outside the geographic range where it had previously been reported. Acceptable evidence of a range contraction was defined as systematic documentation of the absence of a vector species over a substantial period of time (i.e., multiple years) in surveys, locations, and times of the year in which it was previously found regularly. Studies that only presented a modeled projection of vector occurrence without primary empirical data were excluded. Reviews and evidence syntheses were excluded, but the sources they included were checked for possible inclusion.

Study titles and abstracts were screened by two independent reviewers for inclusion or exclusion, with a third reviewer serving to resolve any conflicts. The 299 papers that passed the first screen were then read in full by two reviewers (Fig. S1). From these, the reviewers extracted the following information from the studies that met the inclusion criteria (n = 178): species name(s); study geographic location(s), with latitude and longitude points if available; study date(s); reference period date(s); vector collection methods; whether the study reports a range expansion, contraction, or both; distance or velocity of range movement; a narrative summary of the asserted causes of the range shift; a summary of the evidence used in attributing the causes. The reference period is defined as the historical period to which the species range is being compared, if stated.

We summarized data using R version 4.1.1, quantifying the prevalence of different study characteristics using counts and percentages. To visualize study locations, we recorded the latitude and longitude of one representative study site per paper. If coordinates were not provided, we georeferenced the study location based on descriptions (e.g., city, region) and assigned coordinates to the geographic center of the reported area.

## Results and Discussion

### Overview of studies

We identified 2,356 publications in our search and synthesized 178 studies that met our criteria. These studies encompassed 118 unique mosquito species that vector human diseases. The majority focused on *Aedes* species, particularly invasive mosquitoes such as *Aedes albopictus*, *Aedes aegypti*, and *Aedes japonicus* (Fig. 1A). *Anopheles* species also featured prominently, representing roughly a fifth of the studies, while *Culex* species accounted for 14%, and a small portion examined other genera (6%). Survey data for these studies spanned a large temporal range, using data from as far back as 1898, but most were conducted relatively recently, with a median sampling year of 2004 (Fig. S2). The studies were published between 1988 and 2023, with most appearing within the last decade (Fig. 1B). Geographically, the studies have a global reach, covering all continents except Antarctica, where mosquito vectors are not known to exist (Fig. 1C). North America and Europe dominate the study locations, with dense clusters in the Eastern United States and Western Europe. Despite their relatively small land area, islands account for a considerable fraction of studies (12%), likely because colonization events on islands represent clear instances of range expansion.

**Figure 1.**
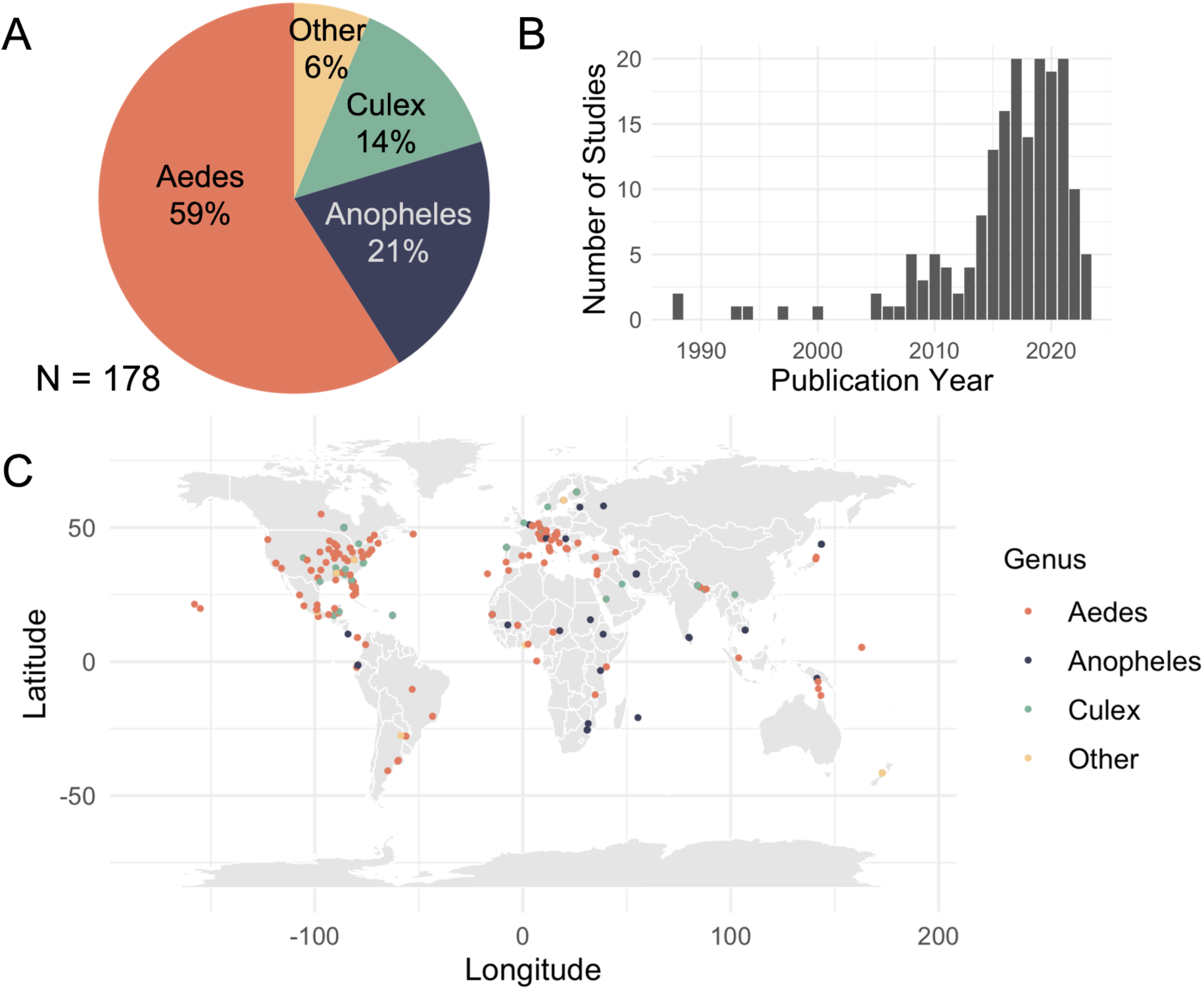
(A) Fraction of studies by mosquito species investigated, with most common three labeled by genus and all others labeled as “Other”. (B) Publication year. (C) Locations where studies were conducted.

Although climate change was often hypothesized as a contributing factor to mosquito range shifts, it was rarely tested through statistical analyses. Over a third of studies in this review (34%) discussed climate change as a potential contributing factor to changes in species distributions. However, the linkage between climate change and range shifts was typically described briefly and qualitatively, rather than through rigorous statistical analysis. Of the studies that discussed climate change, only 9% applied some form of quantitative analysis to test its impact on mosquito range shifts. This included a variety of approaches, such as generalized linear models (GLMs) that incorporate temperature variables (e.g., Campos 2011, Gouagna et al. 2011, Dhimal et al. 2014a, Mogi and Tuno 2014), though often without considering long-term temperature trends. Several studies performed spatial analyses and incorporated temperature variables into environmental niche models or climate envelope models (Caminade et al. 2012, Hopperstad and Reiskind 2016, Kulkarni et al. 2016, Salahi-Moghaddam et al. 2017, Trajer et al. 2017, Rubio et al. 2020). These studies used temperature data as a predictor in their models but generally did not disentangle the role of anthropogenic climate change from natural temperature variability. Another approach involved comparing recent mosquito range expansions to static isotherms, e.g., annual or cold month temperature thresholds (Armstrong et al. 2017, Zanotti et al. 2015). These studies assessed whether newly established populations fall within previously identified thermal limits, but did not analyze whether climate change was responsible for recent expansions, nor did they test whether isotherms themselves have shifted over time. Notably, all studies that provided statistical evidence for climate-driven expansions focused on *Aedes* or *Anopheles spp.*, with no comparable analyses for *Culex*. Although climate change was hypothesized as a driver for all three genera, the absence of statistical testing for *Culex* highlights an important gap in understanding their climate responses.

### Case studies with compelling evidence for climate change impacts

Several studies provided clear evidence of warming-mediated mosquito range expansions using a variety of methods for data collection and analysis. A common goal among these studies was to establish both the occurrence of warming in a certain area and estimate the relationship between mosquito occurrence and temperature. We highlight three representative case studies that provided compelling evidence, as a means of outlining methodological approaches for attributing mosquito range shifts to climate change.

One example is the southern expansion of *Ae. aegypti* in South America. Carbajo et al. (2019) examined the change in distribution of *Ae. aegypti* in Argentina by combining georeferenced public records spanning 1986 to 2017 with field surveys conducted in 14 cemeteries in 2017. They fit generalized linear mixed models that accounted for environmental variables such as mean annual air temperature, precipitation, altitude, and human population, to current and historic data and demonstrated a significant association between temperature and the mosquito’s distribution. A follow-up by Rubio et al. (2020) built on this work by conducting active field searches through the region in 2019, ultimately revealing a southwestern expansion to new cities. This expansion aligned with warming trends observed in the region and the future predictions made by Carbajo et al. (2019). However, a limitation of this case study is that it remains unclear whether the new southernmost locations were surveyed during earlier periods when temperatures were unsuitable for mosquito survival—meaning that a true absence record is lacking.

A second example is a study conducted by Mogi and Tuno (2014) on the northern limit of *Ae. albopictus* in Japan. Using long-term surveillance records since 1993, the researchers matched temperature data to the year the vector was sampled and applied a thermal suitability index to assess habitat suitability. Their analysis identified that both insufficient summer warmth and severe winter cold could limit *Ae. albopictus* presence. Although range expansion began in certain localities before rapid warming, the study found that warming has facilitated further northward expansion. This evidence suggests that warmer winters in particular have played a critical role in enabling this species to extend its range into previously unsuitable areas. The authors suggest that future studies fit multifactor climatic-niche models based on distribution data or physiological requirements of the mosquito.

A third example is Pedrosa (2021), which documented the expansion of *Aedes aegypti* and *Aedes albopictus* into high-elevation tropical cities in southeastern Brazil from 2009 to 2019, showing that their persistence depended on rising winter temperatures. Field surveys across multiple montane urban areas in 2009 and 2011 revealed stable populations above 1,000 meters, where these species were previously rare. Regression analyses indicated that while summer conditions have long been suitable, recent increases in winter minima have allowed populations to establish year-round. The study underscores how warming winters, rather than summer extremes, can drive mosquito range shifts into higher elevations. Following this expansion, dengue cases in the region increased sharply, with autochthonous infections becoming more frequent over time.

Collectively, these studies include several features that enable compelling attribution of mosquito range expansions to climate change. First, robust evidence requires long-term datasets that capture both historic and current temperature and mosquito occurrence records. Such datasets can capture whether a species was previously limited in range by temperature (i.e., true negatives) but has recently expanded beyond that limit. Additionally, employing statistical analyses—whether through regression models, environmental niche models, or thermal suitability indices—that control for confounding factors like humidity and urbanization is essential to isolate the effect of climate warming. Finally, two of the three case studies (Mogi and Tuno 2014 and Pedrosa 2021) examined specific temperature variables to identify that warming winter temperatures, potentially even more importantly than warming activity-season temperatures, have contributed substantially to *Aedes* spp. range expansions. In all three case studies, expansions of *Ae. aegypti* and *Ae. albopictus* carry immediate risk for transmission of dengue, Zika, chikungunya, and other arboviruses if they were introduced from endemic regions.

### Invasive *Aedes albopictus*, *Aedes aegypti*, and *Aedes japonicus*

Studies involving the invasion of *Aedes albopictus*, *Aedes aegypti*, and/or *Aedes japonicus japonicus* made up more than a third (62) of the total studies. Most of these studies are new reports in areas that have always been thermally suitable and the documented range expansion is representative of the mosquito filling out its fundamental niche. Globally, *Ae. aegypti* frequently spreads via long-distance importation, into areas with pre-existing thermal suitability, while *Ae. albopictus* tends to expand outward along the fringes of its established range distribution (Kraemer et al. 2019). Range expansion of *Ae. japonicus* is largely documented to occur with human movement, tourism, and urban expansion. In higher altitudes, Aditya et al. (2009) and Dhimal et al (2014a, 2015) documented *Ae. aegypti* presence at elevations above 2,000 m in the Himalayas, significantly higher than previous records at lower altitudes (e.g., Kurseong at 900m and Rongtong at 482m). The role of increased tourism and human movement in aiding dispersal, as suggested by these studies, is a key alternative driver of range shifts and will be explored in detail in the following section.

However, some records show expansion of these invasive species northward and upward in elevation that may be due to climate change. Armstrong et al. (2017) studied *Ae. Albopictus* populations across a 20-year span at fixed trap sites in Connecticut. Mosquito abundance increased steadily from 2010 onward, with exceptions in 2013-2014 and 2015-2016—both colder winters. Correlating *Ae. albopictus* presence with a 0°C winter isotherm suggests that milder winters may improve overwintering survival and facilitate range expansion, consistent with the case studies discussed above (Mogi and Tuno 2014 and Pedrosa 2021). In Austria, Bakran-Lebl et al. (2021) observed expansion patterns of *Ae. albopictus,* documenting northward shifts, with hypothesized expansion in other directions. Similarly, Ruiz-Lopez et al. (2016) reported altitudinal increases of *Ae. albopictus* in Colombia, linking these shifts to favorable conditions in higher elevation zones. Thus, while much of the spread of these globally invasive species has been into already suitable areas, the poleward and altitudinal directions of range expansions suggest the potential for climate change as a driver. Given the predominantly observational nature of these studies, future research should incorporate additional statistical analyses that more rigorously assess the role of climate change in these expansions.

### Alternative hypotheses for range expansion

A total of 105 studies (60%) proposed alternative explanations for range expansion, many of which involved interrelated or interacting factors. The most commonly cited driver was dispersal, either through human-assisted movement via commerce and human transit (74 studies) or through extreme weather events (four studies) (Table 1). These mechanisms facilitate long-distance transport and introduce mosquitoes to new regions where conditions may already be suitable for establishment. *Ae. albopictus* was most commonly discussed in the context of human-mediated dispersal with 18 studies hypothesizing that the observed expansion was due to trade of goods, especially tires and bamboo, and 19 studies hypothesizing that the expansion was due to increased human movement through development of roadways, railways, air travel, and maritime shipping. A number of other *Aedes* spp., including *Ae. japanicus, Ae. bahamensis*, *Ae. keoricus,* and *Ae. scapularis*, as well as the container-breeding *Anopheles stephensi*, are also believed to have experienced changes in distribution due to trade and human movement. Notably, all *Aedes* species in these studies are speculated to have spread specifically through the trade of tires and bamboo, whereas *An. stephensi* may have spread through the movement of livestock (Carter et al. 2021) and, in some cases, fishing boats (Gayan Dharmasiri et al. 2017). The evidence for these dispersal pathways was indirect and circumstantial, relying on associations with human activity rather than the tracking of mosquito movement. For instance, Alarcón-Elbal et al. (2014) inferred that *Ae. albopictus* spread through roadways based on the correlation between *Ae. albopictus* abundance and traffic volume, whereas other studies (Cvetkovikj et al. 2020, Fuehrer et al. 2020) drew similar conclusions based on anecdotal evidence that the species was frequently captured close to bus stations or railways. Finally, *Culex* spp. (*Cx. coronator, Cx. nigripalpus*) and *Ae. bahamensis* were thought to have dispersed via wind caused by extreme weather events, such as hurricanes (Pafume et al. 1988, Connelly et al. 2016, Akaratovic and Kiser 2017). However, this was based on previous studies and the observation that adults were found in unexpected locations following storms (Akaratovic et al. 2022). Conclusively identifying the role of human-mediated dispersal may require mark-recapture or landscape genomic approaches (e.g., Marcantonio et al. 2019)

**Table 1.**
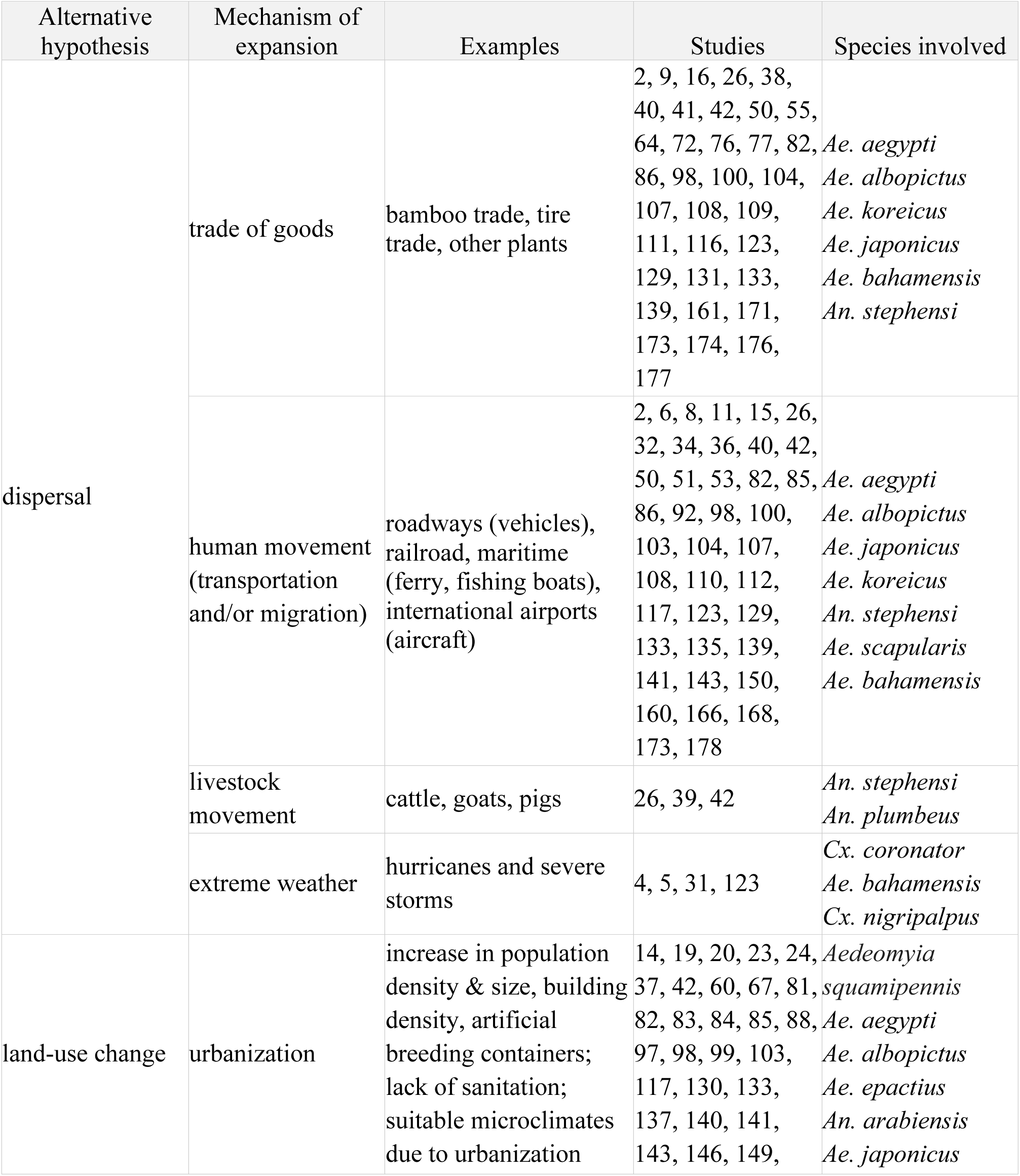
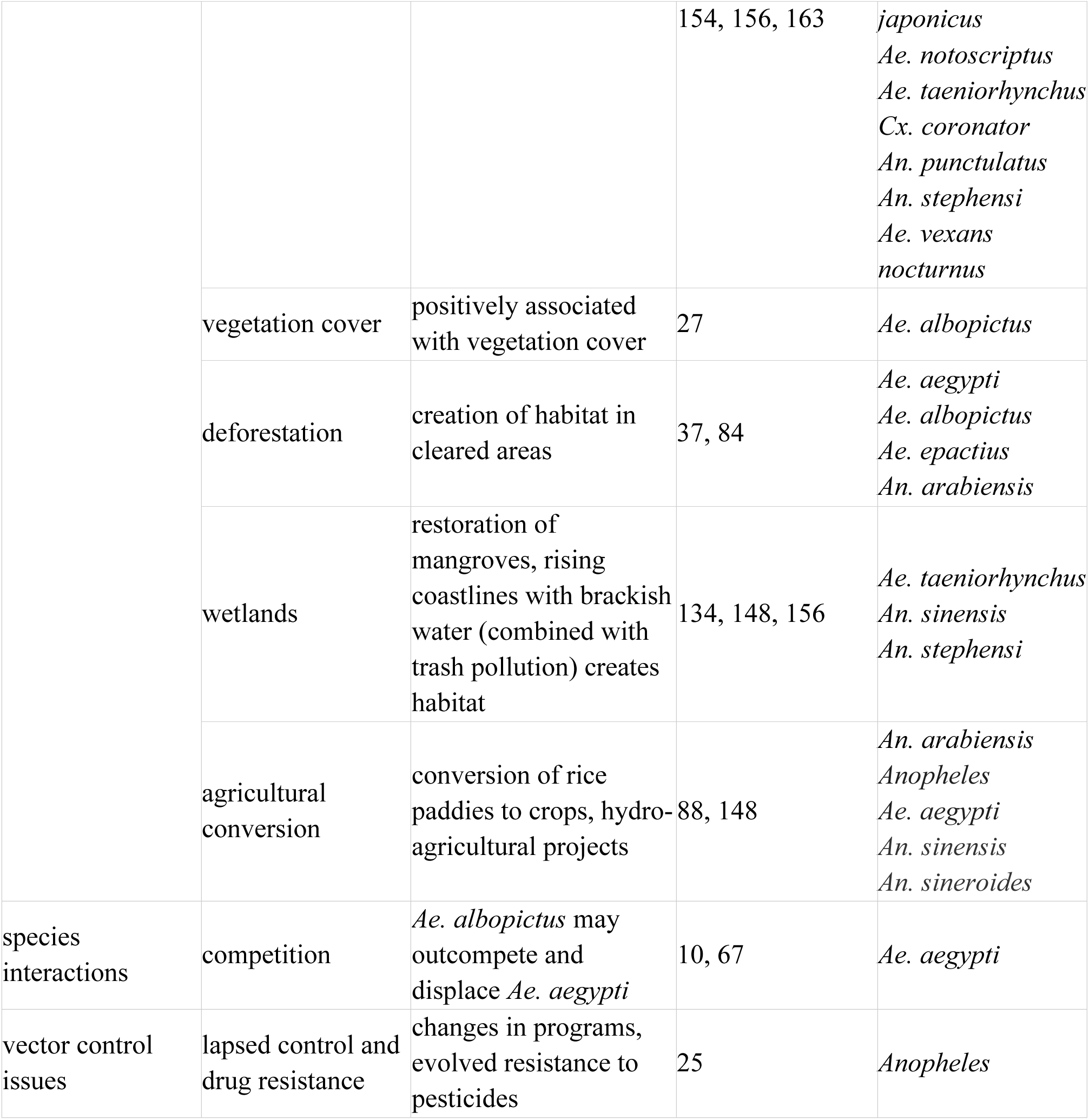
Alternative hypotheses for mosquito range expansion. Studies proposing non-climate explanations for observed range shifts are categorized by hypothesized mechanisms of expansion and species involved.

Land-use change was the next most frequently cited factor potentially driving range expansions, mentioned in 34 studies (19%). Urbanization was the predominant type of land-use change linked to range expansion (31 studies), described in terms of both urban expansion and population growth. As cities expand, they create conditions that favor mosquito establishment, such as an abundance of artificial breeding sites (e.g., plastic bottles, tires), the development of urban heat islands, and inadequate sanitation. *Aedes aegypti* was most often associated with urbanization (16 studies), followed by *Ae. albopictus* (8 studies), other *Aedes* and *Anopheles* species, *Cx. coronator*, and *Aedeomyia squamipennis* (Table 1). Other forms of land-use change implicated in range expansions included deforestation, wetland modification, and agricultural expansion or conversion. While most hypotheses involving land-use change were based on anecdotal evidence or previous studies, two studies provided quantitative evidence. Prioteasa and colleagues (2015) found that population size, a frequently used proxy for urbanization, was a strong predictor of *Ae. albopictus* presence. Qualls et al. (2022) incorporated land-use/land-cover variables and a long-term surveillance dataset in their analysis of *Ae. taeniorhynchus* distribution changes, finding that wetland restoration expanded mangrove swamp habitat, leading to a broader distribution of the species.

Beyond dispersal and land-use change, researchers also proposed that changes in species ranges could be due to competition between *Ae. aegypti* and *Ae. albopictus* (Bennet et al. 2021, Hopperstad and Reiskind 2016) and lapses in vector control and/or development of insecticide resistance (Carlson et al. 2023a). Competition was believed to have displaced *Ae. aegypti*, as lab and field research indicates that *Ae. albopictus* can outcompete *Ae. aegypti* both through dominance of *Ae. albopictus* larvae in shared resource consumption and breeding interference (Braks et al. 2004, Gilotra et al, 1967, Lounibos and Juliano 2018).

### Range contractions and reasons for them

None of the studies we screened provided clear evidence of vector range contractions linked to climate warming. In total, 8 studies (4%) described observed contractions in vector distributions or reductions in abundance (e.g., through comparisons of historical and current surveillance records) (Gouagna et al. 2011, Dekoninck et al. 2013, Kulkarni et al. 2016, Lemine et al. 2017, Salahi-Moghaddam et al. 2017, Bennett et al. 2021, Sawabe et al. 2021, Qualls et al. 2022). Of these, one study did note a concurrent trend of increasing temperatures. Kulkarni et al. (2016) documented a contraction in the distribution of *Anopheles arabiensis* in lower altitude regions of Tanzania (and an expansion at higher altitude regions) between the 2001-2004 and 2014-2015 sampling periods. Further, annual mean temperature was found to have increased across this time period, and found to be a relatively strong predictor of shifting species distributions across the study region. However, this appears to be driven by the role of temperature in promoting vector suitability at the higher altitudes, as noted by the authors, “hot season temperature was associated with increasing suitability for *An. arabiensis* occurrence”, and the contractions at lower altitudes are described as being driven primarily by urbanization. The remaining 7 studies noting contractions either found no effect of temperature (Salahi-Moghaddam et al. 2017) and/or did not investigate effects of temperature but described the drivers being competitive displacement by another mosquito species (Bennett et al. 2021), precipitation (Gouagna et al. 2011), and/or land use change (Lemine et al. 2017, Bennett et al. 2021, Sawabe et al. 2021, Qualls et al. 2022). Whether the lack of observed warming-related range contractions is due to methodological limitations and/or sampling effort (i.e., challenges in observing the absence of a species), a true lack of contractions (e.g., due to evolutionary or plastic adaptive responses), potentially lagged impacts of warming (e.g, ‘extinction debt’), or a lack of recognition that climate change may lead to temperatures above species’ upper thermal limits remains unclear (Couper et al. 2021, Couper et al. 2024).

### The speed of range expansion

There were 16 studies (9%) that provided sufficient information to estimate the speed of mosquito range expansion. Most of the studies that reported poleward expansions observed rates between 2.5 and 13.6 km/year (Bakran-Lebl et al. 2021, Lemine et al. 2017, Mogi and Tuno 2014, Carlson et al. 2023a). Kraemer et al. (2019) was a notable outlier, reporting continent-scale expansions of *Ae. albopictus* and *Ae. aegypti* at rates of 60, 150, and 250 km/year, roughly an order of magnitude higher than the other studies. Additionally, eight studies reported expansions in directions other than poleward and are likely unrelated to warming climates. In terms of altitudinal expansion, three studies (Romiti et al. 2022, Carlson et al. 2023a, Pinault and Hunter 2011) found that mosquito species are moving into higher elevations at rates ranging from 6.5 to 17 m/year. Both the poleward and elevational expansion rates documented for mosquitoes are substantially higher than those found in other taxa, where a meta-analysis by Chen et al. (2011) reported median rates of 1.7 km/year for latitudinal shifts and 1.1 m/year for altitudinal shifts. A key limitation of these speed estimates is that most were derived from a single distance measurement between the range edge in two surveys separated in time by a few years (Klobučar et al. 2019) to several decades (Lemine et al. 2017). However, some studies have taken a more robust approach, using multiple time points (Bakran-Lebl et al. 2021, Tavecchia et al. 2017) or regressions on long-term datasets (Kraemer et al. 2019, Carlson et al. 2023a). Another limitation is that the observed poleward and altitudinal expansions are not necessarily driven by climate change but could occur as species fill in areas already within their fundamental climatic niche, as several studies highlighted the role of long-distance importation (Kraemer et al. 2019) and the "leapfrog" nature of mosquito dispersal (Tavecchia et al. 2017). Furthermore, it is important to note that warming patterns are not uniform, with subtropical regions experiencing faster rates of warming and higher thermal extremes than the tropics. These nuanced changes in temperature regimes could significantly impact mosquito population dynamics and range expansion, beyond simple latitudinal trends.

### Consequences for disease transmission

The expansion of mosquito ranges has significant consequences for the spread of vector-borne diseases, as novel occurrences of both pathogens and their vectors can emerge in previously unaffected areas. For instance, in both the mountains of Nepal (Dhimal et al. 2014a, 2014b, 2015) and eastern Brazil (Pedrosa 2021), dengue cases were reported in high elevation areas where *Aedes* mosquitoes, traditionally not found in these regions, had recently established. Similarly, in Argentina, there has been a notable increase in both the incidence and geographic expansion of dengue, where warming temperatures have been shown to influence the dynamics of dengue outbreaks (López et al. 2023). These case studies highlight the growing concern over the spread of *Aedes*-vectored diseases, such as dengue, Zika, and chikungunya, in regions previously not considered at risk. Other studies have shown how past and future climate change is affecting the distribution of *Anopheles*, contributing to shifts in malaria transmission dynamics in Africa (Ryan et al. 2015, Peterson 2009; Carlson et al. 2023b). As many different mosquito vectors continue to expand their ranges, we may witness further geographic spread of diseases like malaria and dengue, presenting new challenges for public health systems.

### Open questions and next steps

#### 1. How can we more rigorously attribute range shifts to climate warming?

Although temperature is frequently mentioned as a driver, few studies directly connect observed warming trends to mosquito expansions. Long-term monitoring datasets (e.g., those in Armstrong et al. 2017, Stone et al. 2020) could shed light on this question if paired with formal attribution methods (e.g., comparing distribution changes to shifting isotherms, species distribution models with counterfactual climate scenarios (Glidden et al. 2024). The increasing availability of remotely sensed environmental data and automated mosquito species detection tools (e.g., González et al. 2022), offers new opportunities to address this question at broader spatial and temporal scales.

#### 2. Are mosquitoes filling out their existing niche or expanding into newly thermally suitable regions?

Many expansions occur in areas that were already climatically suitable, suggesting that lack of prior detection—or recent introductions via trade and travel—may explain many of the newly reported populations. Thus, there is a need to untangle climate-driven range expansion from the filling of unoccupied suitable habitat. Climate change alters mosquito ranges through mechanisms like warmer winters increasing adult survival. However, many studies rely on annual mean temperatures, which may not be the primary drivers of selection and population dynamics.

#### 3. Why are warm-edge contractions so rarely observed?

No clear examples attribute lower-range contractions to overheating or reduced fitness. Is this due to behavioral or evolutionary adaptive responses, limited sampling effort, or extinction debts, in which species declining due to locally unsuitable conditions have not yet gone locally extinct? Understanding how mosquitoes adapt to high-temperature limits remains an important research frontier, particularly as upper thermal limit constraints can be difficult to detect in species distribution models (SDMs) or observational field data (Athni et al. 2024).

#### 4. How do vector control and land-use change interact with climate factors?

Shifts in distribution may be confounded by local or regional vector control measures, as well as changing urban landscapes. Future studies must document these interventions to better isolate the specific contribution of warming from other anthropogenic drivers that can indirectly affect microclimates.

#### 5. What are the public health implications of shifts in mosquito abundance and phenology?

Most investigations emphasize the presence or absence of species in newly invaded regions, rather than changes in abundance, seasonality, or vector competence. Addressing these gaps is essential for predicting and managing emerging disease risks under continued climate change.

## Conclusions

Mosquito species of public health concern are rapidly expanding their ranges, at rates that are higher than many other taxa. These range expansions are likely driven by a combination of anthropogenic factors including climate change, land use change, and human movement. Despite clear expectations from thermal biology that climate warming will expand mosquito species distributions at their cool range limits, and evidence that many taxa, including *Aedes*, *Anopheles*, and *Culex* spp., are expanding poleward and upward, rigorous evidence of climate warming-driven range expansions remains limited to several compelling case studies. These studies suggest an important role of warmer temperatures year-round and especially in the winter in allowing *Aedes* mosquitoes to establish in more temperate regions. Concurrent range expansions (i.e., invasions) into existing thermally suitable areas present a challenge for quantifying and attributing the role of anthropogenic climate change. Although empirical and theoretical work suggests that climate warming could drive warm-edge range contractions, there is no compelling evidence of this to date for any mosquito species. Overall, quantifying the extent of mosquito range expansions due to climate change remains a major research gap that requires coordinated, long-term vector surveillance in areas of emerging climate suitability. Such efforts could jumpstart public health responses in areas facing novel risk of mosquito-borne disease outbreaks that previously lacked vectors, including highland and temperate regions of Asia, the Americas, Europe, Africa, and Australia and Oceania.

## Acknowledgements

KL was supported by the NSF Postdoctoral Research Fellowships in Biology Program under Award No. 2208947. LIC was supported by the NSF Postdoctoral Research Fellowships in Biology Program under Award No. 2410030. CKG was supported by a Stanford Institute of Human-centered Artificial Intelligence Postdoctoral Fellow and the Doerr School of Sustainability Center for Human and Planetary Health. IOD was funded by the National Institutes of Health (R35GM133439). EAM was supported by the National Institutes of Health (R35GM133439, R01AI168097, R01AI102918), the National Science Foundation (DEB-2011147, with Fogarty International Center), and the Stanford Center for Innovation in Global Health, King Center on Global Development, and Woods Institute for the Environment.

