## Supplement for "A systematic review of climate-change driven range shifts in mosquito vectors"

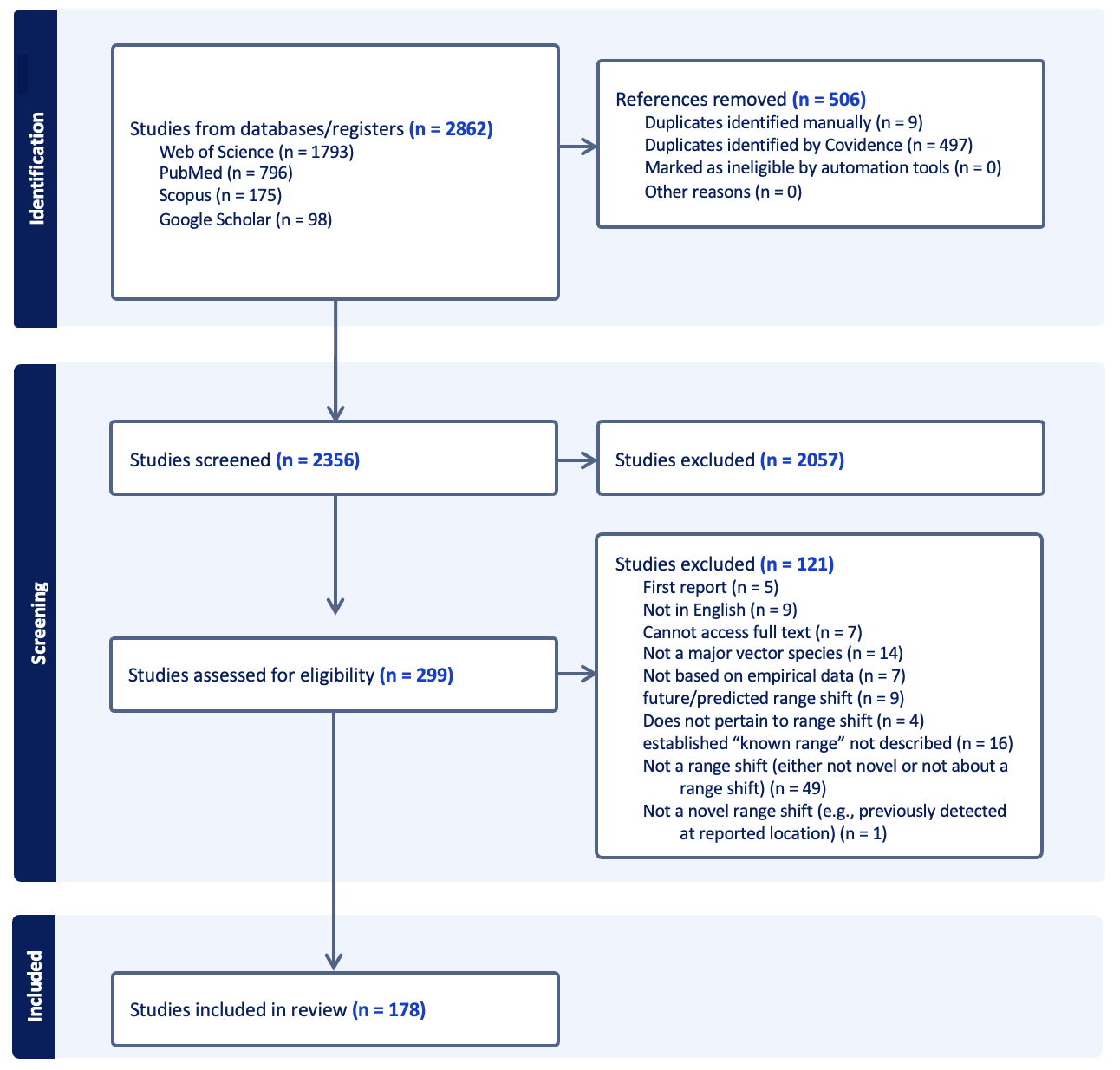

**Figure S1.** Flow chart of studies processed in the systematic review.

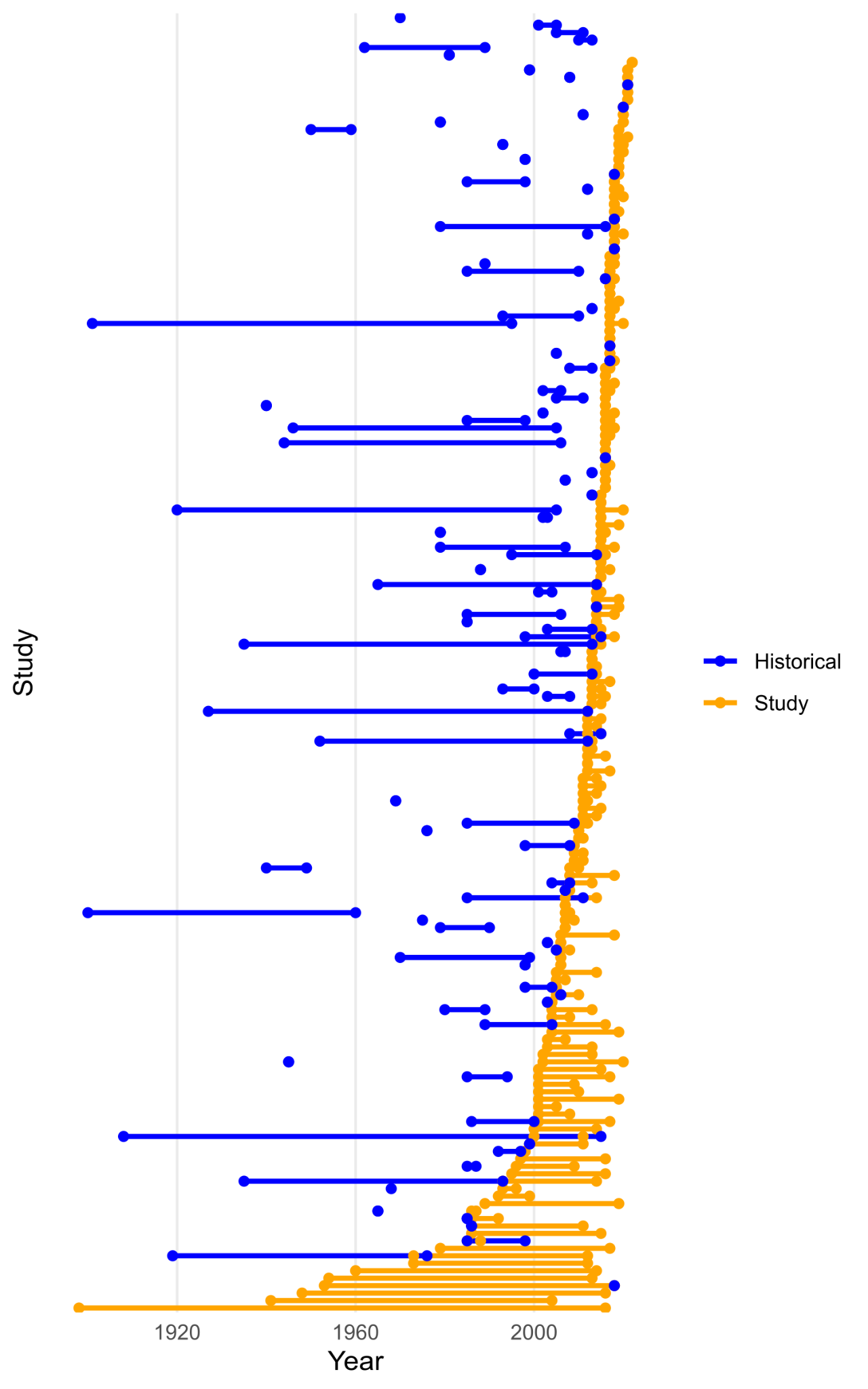

**Figure S2.** Time periods of mosquito surveys are short and recent. Yellow indicates the years of surveys conducted or survey data analyzed within the study, while blue represents historical surveys used as reference points. Studies are arranged chronologically by the first yellow year.
